# Tree-based QTL mapping with expected local genetic relatedness matrices

**DOI:** 10.1101/2023.04.07.536093

**Authors:** Vivian Link, Joshua G. Schraiber, Caoqi Fan, Bryan Dinh, Nicholas Mancuso, Charleston W.K. Chiang, Michael D. Edge

## Abstract

Understanding the genetic basis of complex phenotypes is a central pursuit of genetics. Genome-wide Association Studies (GWAS) are a powerful way to find genetic loci associated with phenotypes. GWAS are widely and successfully used, but they face challenges related to the fact that variants are tested for association with a phenotype independently, whereas in reality variants at different sites are correlated because of their shared evolutionary history. One way to model this shared history is through the ancestral recombination graph (ARG), which encodes a series of local coalescent trees. Recent computational and methodological breakthroughs have made it feasible to estimate approximate ARGs from large-scale samples. Here, we explore the potential of an ARG-based approach to quantitative-trait locus (QTL) mapping, echoing existing variance-components approaches. We propose a framework that relies on the conditional expectation of a local genetic relatedness matrix given the ARG (local eGRM). Simulations show that our method is especially beneficial for finding QTLs in the presence of allelic heterogeneity. By framing QTL mapping in terms of the estimated ARG, we can also facilitate the detection of QTLs in understudied populations. We use local eGRM to identify a large-effect BMI locus, the *CREBRF* gene, in a sample of Native Hawaiians in which it was not previously detectable by GWAS because of a lack of population-specific imputation resources. Our investigations can provide intuition about the benefits of using estimated ARGs in population- and statistical-genetic methods in general.

## Introduction

Identifying trait-associated genetic loci is one of the central aims of genetics. Over the past several decades, a range of approaches—prominently including linkage mapping and genome-wide association studies (GWAS)—appeared to fill this need (Balding et al., 2019). In humans, GWAS has become a tremendous research enterprise, with millions of study participants enrolled and hundreds of thousands of trait-associated variants identified (Visscher et al., 2017).

For decades, geneticists have noted the usefulness of tree-based structures for describing genetic variation and for characterizing the genealogical and evolutionary processes that create genetic variation. At a single non-recombining locus, a tree called a gene genealogy describes the shared ancestry of individual copies of the locus (Rosenberg and Nordborg, 2002). For entire genomes or genomic regions in which recombination events occurred in the history of the sample, one can represent the sample’s shared ancestry via an ancestral recombination graph (ARG) that encodes the sequence of "local" or "marginal" trees along the genome (Griffiths and Marjoram, 1996), with recombination events as the source of differences in topology between neighboring trees. The ARG encodes all mutation, recombination, and shared ancestry events in the history of a sample of genomes.

Tree-based approaches to quantitative trait locus (QTL) mapping—in which a trait is tested for association with a tree or set of trees describing genetic variation in a region—have been proposed several times and shown to provide some advantages (Templeton et al., 1987; McPeek and Strahs, 1999; Larribe et al., 2002; Morris et al., 2002; Zöllner and Pritchard, 2005; Minichiello and Durbin, 2006; Mailund et al., 2006; Tachmazidou et al., 2007; Kimmel et al., 2008; Wu, 2008; Besenbacher et al., 2009; Zhang et al., 2012; Burkett et al., 2013; Thompson and Kubatko, 2013; Thompson et al., 2016), as have approaches to haplotype-based mapping that leverage awareness of tree-like relatedness patterns among sets of haplotypes (Liu et al., 2001; Morris, 2005; Selle et al., 2021). At the same time, explicitly tree-based approaches have until recently been limited by difficulties in estimating locus-level trees at scale. Further, the dominance of meta-analysis in GWAS (Cantor et al., 2010) and other methods based on summary statistics has meant that individual-level genetic data are often not available to data analysts, precluding most tree-based approaches.

In principle, tree-based approaches have the potential to address three long-standing difficulties of GWAS. First, GWAS entails a huge number of statistical tests and requires a substantial correction for multiple testing as a result (Pe’er et al., 2008). Many of these tests are correlated or redundant because the variants tested occur on the same or very similar underlying gene-genealogical trees. Testing the trees themselves may allow for fewer tests.

Second, GWAS is known to be prone to miss trait-associated genetic loci characterized by allelic heterogeneity, in which multiple nearby causal variants affect a trait of interest (Platt et al., 2010; Flister et al., 2013; Korte and Farlow, 2013; Hormozdiari et al., 2017). Under allelic heterogeneity, causal alleles with opposing effects on a trait might be associated with the same marker allele, diminishing the association signal at the marker. Allelic heterogeneity is not rare, appearing in many Mendelian loci identified during the linkage era (Terwilliger and Weiss, 1998)—linkage mapping is robust to allelic heterogeneity—and estimated recently to occur at a substantial fraction of complex trait loci (Hormozdiari et al., 2017) and expression QTLs (Jansen et al., 2017; Abell et al., 2022). Tree-based approaches, by focusing on local relatedness of haplotypes in the sample, can offer the same robustness to allelic heterogeneity as linkage analysis.

Third, modern GWAS is fueled by imputation, in which a reference sample is fully sequenced, and then study samples that are more sparsely genotyped have their missing genotypes imputed statistically (Marchini and Howie, 2010; Das et al., 2018). The imputed genotypes can then be tested for association with the trait of interest. The success of modern imputation approaches is made possible by the fact that genetic variation is structured locally in a tree-like way (Stephens and Scheet, 2005; Edge et al., 2013). At the same time, imputation is most successful if the reference and study samples are closely related (Huang et al., 2009; Jewett et al., 2012; Lin et al., 2020), and closely related reference samples are not always available. Testing the tree structures that underlie imputation may offer a more direct approach to identifying QTLs that could circumvent the need for closely related reference samples.

Due to advances in ARG estimation, it may now be possible to apply tree-based methods at sufficient scale to detect QTLs. Although estimation of the ARG is extremely difficult, approximate estimation procedures that operate on single-nucleotide polymorphism (SNP) array data and scale to thousands of samples have emerged in the last few years (Kelleher et al., 2019; Speidel et al., 2019; Zhang et al., 2021; Wohns et al., 2022). Further, for researchers studying QTLs in humans, the emergence of large biobanks has meant that individual researchers or research teams have access to individual-level genetic data in sample sizes that might allow the identification of trait-associated loci.

Here, we present a tree-based approach to QTL mapping (Figure 1). We build on a recently proposed representation of tree-based relatedness, the expectated genetic relatedness matrix, or eGRM (Fan et al., 2022, see Figure 1B here), independently identified by Zhang and colleagues (2021) as the ARG-GRM and in a phylogenetic context by Wang and colleagues (2021) as the expected genetic similarity matrix. (And see McVean (2009, eq. 10) for a similar computation.) Genetic relatedness matrices (GRMs) are used in a wide array of statistical genetic tasks, including adjusting for population stratification and estimation of heritability (Speed and Balding, 2015). Given an ARG encoding the history of a sample, the eGRM is the expectation of the GRM assuming that mutations are placed on the ARG as a Poisson process. In general, we can compute a tree-based analog of any statistic computed from genetic variation by taking its expectation given the ARG (Ralph et al., 2020).

**Figure 1:**
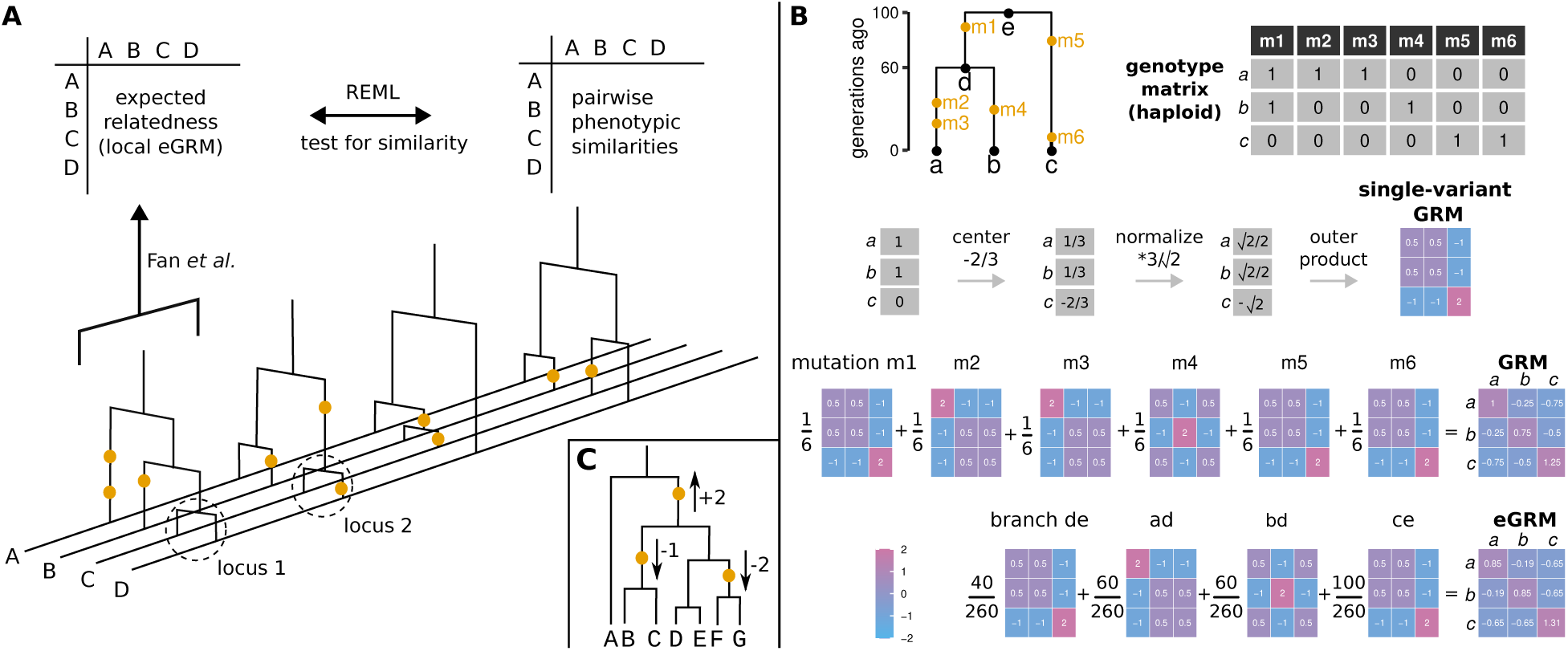
A) Local eGRM framework. Schematic of the marginal trees of an ARG of 4 haploid invdividuals, A-D. The gold circles on the trees correspond to mutations. The clade marked with the dotted circles is identical at loci 1 and 2, and the mutation at locus 2 is informative about the branch length at locus 1. One or more marginal trees are used to calculate a local eGRM using the method described in Fan et al. (2022). This matrix is then tested for association with the phenotypes using Restricted Maximum Likelihood (REML). **B) Computing the eGRM**. This panel is redrawn from Fan et al. (2022). A genome-wide genetic relatedness matrix (GRM) can be viewed as an average of single-locus GRMs for every genotyped locus. An expected GRM (eGRM) can be obtained as a weighted average of single-locus GRMs defined by each branch in the ARG, where the weights are proportional to the expected number of mutations falling on each branch. **C) Allelic heterogeneity**. One marginal tree of an ARG with three causal mutations with opposing phenotypic effects.

Our procedure is to test eGRMs built from local segments of the ARG for concordance with a phenotype using a random-effects model fit by restricted maximum likelihood (REML). Loosely, the test is sensitive to cases in which individuals who are more closely related in some local segment of the genome are likely to be more similar on the phenotype. This approach is essentially a tree-based version of previous methods to test local GRMs for concordance with a phenotype, which have been framed variously as QTL mapping approaches (Wang et al., 2013; Sasaki et al., 2015) or local heritability estimation (Nagamine et al., 2012; Uemoto et al., 2013; Gusev et al., 2013; Caballero et al., 2015). Using tree sequences estimated by `Relate` (Speidel et al., 2019), we test our approach in simulations of varying degrees of allelic heterogeneity. We also use our approach to analyze data from a sample of Native Hawaiians from near the *CREBRF* gene, in which the lack of a population-specific reference panel has previously precluded the detection by GWAS of a known large-effect polymorphism (Minster et al., 2016; Lin et al., 2020).

## Methods

### Characterizing local relatedness

The key to our approach is a matrix **`A`**, called a local genetic relatedness matrix (local GRM), that characterizes the relatedness of individuals in a local region to be tested as a candidate QTL. Classically, a local GRM is calculated based on the observed variants in a a window (e.g. Yang et al., 2010). Our method is instead based on using the expectation of a local GRM (local eGRM) given an estimated ancestral recombination graph (ARG).

Fig. 1 describes the main ideas of our framework: in each window, we calculate one genetic relatedness matrix using the ARG’s marginal trees in that window, and then use REML (Restricted Maximum Likelihood) to test whether the local genetic relatedness explains phenotypic similarities in the sample. One advantage of using the ARG to describe genetic relatedness is that information from neighboring trees is naturally shared. To illustrate this idea, the dotted circles in Fig. 1A show a clade that exists in trees 1 and 2, with a mutation at locus 2 that differentiates tips C and D. Though the mutation is at marginal tree 2, its presence is informative about the branch lengths at marginal tree 1, since the relevant subtree is identical in marginal trees 1 and 2.

Figure 1B describes the method we use to calculate the pairwise expected relatedness matrix (eGRM), developed by Fan et al. (2022), and also how the genetic relatedness matrix is conventionally calculated. Figure 1C shows an example of allelic heterogeneity: multiple causal alleles are in close linkage (e.g. on the same marginal tree of the ARG), so tag SNPs will be linked to several causal alleles with opposing effects. If the causal variants are themselves untyped, this can lead to association signals interfering or even cancelling each other out at the typed variant. Even if the causal variants are typed, association power will be improved if they are tested for association jointly rather than separately.

#### Local GRM

We compute local GRMs from biallelic variants in the local window to be tested as a QTL (e.g. Yang et al., 2010). The entry relating individuals *`i`* and *`j`* in the GRM can be written as

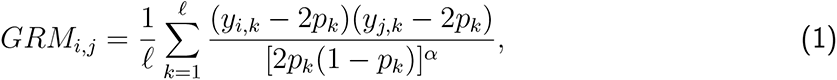

where *`k`* in an index over the sites considered, *`ℓ`* is the total number of sites considered, *`y`_i,k_* is the number of focal alleles carried by individual *`i`*, and *`p`_k_* is the sample frequency of the focal allele (Speed and Balding, 2015). The constant *`α`* determines the relative emphasis placed on rarer variants in computing relatedness estimates, with larger values giving greater weight to rarer variants. In this paper, we use *`α`* `= 1`. With *`α`* `= 1`, the local GRM is a covariance matrix of m<u>ean-cente</u>red, standardized genotype counts among individuals, where the standardization is by 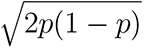, the standard deviation of the genotype under Hardy–Weinberg equilibrium.

#### Local eGRM

Fan et al. (2022) compute a genome-wide global eGRM, which is the expectation of the genetic relatedness matrix (GRM) described by eq. 1 (with *`α`* `= 1`) conditional on the ARG, assuming that infinite-sites mutations are placed on the ARG as a Poisson process. The global eGRM can be computed as a weighted sum of single-locus GRMs implied by each branch in the ARG. Specifically, each branch in the ARG defines a clade of tips that descend from the branch. A mutation on that branch would be inherited by all these tips and so would define a single-locus GRM following eq. 1. The eGRM is equal to the weighted average of all such branch-wise GRMs, with weights per branch proportional to a product *`µ`*(*`b`*)*`l`*(*`b`*)*`t`*(*`b`*), where *`µ`*(*`b`*) is the mutation rate on the branch, *`l`*(*`b`*) is the length of the genomic region spanned by the branch, and *`t(b)`* is the length of the branch (i.e. the time in the tree that the branch exists). Here, as in Fan et al. (2022), we assume that the mutation rates are the same on all branches.

We compute the local eGRM for genomic regions of a pre-defined size. The local eGRM for one tree is a weighted sum over the tree’s branches. In order to calculate the local eGRM for a genomic window, we first calculate the local eGRM for all trees whose genomic intervals overlap the window, and then take a weighted average of these matrices, where the weights correspond to the fraction of the window covered by each tree’s genomic interval. This approach to computation is redundant because many branches exist across multiple marginal trees, and can be ameliorated in principle via an approach that records unique branches only once (Ralph et al., 2020). We did not pursue this solution because of our decision to work with `Relate` trees, which do not preserve branch lengths exactly between neighboring marginal trees.

We computed local eGRMs using `egrm` software (Fan et al., 2022).

### The variance-components model

Let **`y`** be quantitative phenotypes for *`n`* individuals, **`X`** be an *`n`* `x` *`k`* design matrix of covariates, and *`β`* the covariates’ regression coefficients. The *`k`* covariates may include nuisance variables and potentially confounding factors, such as age, sex, and descriptions of population structure or global relatedness. Additionally, let **`I`***_n_* the *`n`* `x` *`n`* identity matrix, and *`σ`*^2^ the variance of environmental noise. Given a GRM **`A`** of dimensions *`n`* `x` *`n`* representing relatedness among individuals in a local segment of the genome, we model the phenotypic variation in the sample by

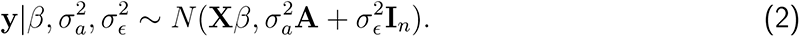

We estimate *`β`*, 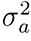, and 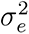 with Restricted Maximum Likelihood (REML). We identify a QTL if the parameter 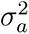 is significantly different from 0, and can further estimate the local heritability as 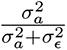.

To understand the difference between using a GRM based on observed variants compared with an expectation conditional on an estimated ARG, note that when **`A`** is based on observed variants, the model in eq. 2 is equivalent to one in which the typed sites in the window receive random, uncorrelated effect sizes with expectation 0 and variance proportional to `1`*`/`*`(2`*`p`*`(1` *− `p`*))*^α^* (Lynch and Walsh, 1998; Goddard et al., 2019) that contribute additively to the trait. That is, *`y`* = **X***`β`* + **Zu** + **e**, with **`X`** an *`n` × `k`* design matrix of covariates with fixed effects *`β`* (*`k` ×* `1`), **`Z`** an *`n` × `ℓ`* matrix of genotypes at the sites considered with random effects **`u`** (*`ℓ` ×* `1`), and **`e`** an *`n` ×* `1` vector of random, uncorrelated environmental effects.

On the other hand, if **`A`** is computed by taking an expectation over an ARG, the model in eq. 2 is equivalent to one in which each branch of the ARG incorporated in **`A`** receives a random, uncorrelated effect size with expectation 0 and variance proportional to *`µ`*(*`b`*)*`l`*(*`b`*)*`t`*(*`b`*)*`/`*`(2`*`p`*`(1`*−`p`*))*^α^*, where *`p`* is the proportion of tips that descend from a branch in the relevant span of the genome, and again *`µ`*(*`b`*) is the mutation rate on the branch, *`l`*(*`b`*) is the length of the genomic region spanned by the branch, and *`t`*(*`b`*) is the length of the branch.

### Simulating genealogy and genotypes

#### One population

For most of our simulations, we simulated ARGs using `stdpopsim` (Adrion et al., 2020; Lauterbur et al., 2022, version 0.1.2) using the Python API. We simulated chromosome 1 for 2000 haploid individuals of African ancestry using the "OutOfAfrica_3G09" model and `msprime` (Kelleher et al., 2016, version 1.1.1) and otherwise default parameters. We then extracted the genomic region starting at position 49,000,000 and ending at position 50,000,000. Then, we randomly assigned pairs of haplotypes to 1000 individuals to create diploids.

#### Two populations

Using `msprime` directly, we simulated two samples of 1000 diploids, sampled from two populations that split 10,000 generations ago and each had a diploid *`N`_e_* of 20,000, which corresponds to an *`F`_ST_* of 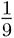 (Slatkin, 1991). We simulated two chromosomes: one “test” chromosome used for association testing of length 100,000bp and and a second chromosome, used to estimate the global eGRM, with length 3,000,000bp. We set the mutation and recombination rates to `10`*^−^*^8^.

Estimating ARGs with **`Relate`** To simulate genotyping array data, we filtered the simulated ARG’s variants by retaining 20% of those with a minor allele frequency of at least 1%. We then used the retained variants to estimate ARGs with `Relate` (Speidel et al., 2019) using parameters ’–mode All’, ’–mutation_rate 1.25e-8’, ’–effectiveN 2000’ and the human recombination map (HapMap phase II, build GRCh37, provided with the `Relate` software). We then converted the output to treeSequence format (Baumdicker et al., 2022) with `Relate`’s tool RelateFileFormats and ’–mode ConvertToTreeSequence’

### Simulating phenotypes

#### Choosing the causal variants

We selected causal variants among those that were not retained in the downsampling scheme described above ("untyped"). In the experiments with one causal variant, the selection was uniformly at random among variants of a predefined frequency. If no branch in the local trees subtended the desired frequency, we chose the nearest possible frequency. In the experiments with allelic heterogeneity, we defined causal regions of different lengths in the center of the ARG and randomly selected a given proportion of untyped variants within the region to be causal.

#### Choosing the effect sizes

We chose the effect size for each variant on the basis of its allele frequency, sampli<u>ng from a</u> normal distribution with expectation 0 and standard deviation inversely proportional to 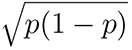 (Speed et al., 2017, LDAK model with *`α`* `= 1`),where *`p`* is the variant’s minor allele frequency. This leads variants with lower allele frequencies to have effects with larger absolute sizes.

In order to obtain the desired local heritability, we added random noise to the phenotypes such that *V_E_* = *V_G_*(1 *− h*^2^)*/h*^2^, where *V_E_* is the phenotypic variance due to environmental effects uncorrelated with genotype, *V_G_*is the phenotypic variance due to genetic effects, and *h*^2^ is the desired local heritability.

### QTL testing

#### local REML

We tested each local relatedness matrix (GRM and eGRM) for association with the phenotypes using `GCTA` (version 1.94.1) (Yang et al., 2011) and its implementation of Restricted Maximum Likelihood (REML) with tag ’–reml’ and providing the local relatedness matrix (tag ’– grm’), the phenotypes (tag ’–pheno’), and running the algorithm for a maximum of 500 iterations. Note that `GCTA` *`p`*-values for random effects in such a model are never larger than `1`*`/`*`2`.

#### GWAS

We tested each typed variant for association with the phenotypes using python’s statsmodels (Seabold and Perktold, 2010) (version 0.13.2) OLS function.

#### ACAT-V

We use function ACAT from R package ACAT (Liu and Xie, 2018, version 0.91) to run ACAT-V on the *p*-values from the GWAS results in a window.

### Correcting for population stratification

We simulated 100 replicates of samples from two populations, as described above. Then, we assigned a random phenotype sampled from *N* `(0`, `1)` to the individuals from the first population, and a phenotype sampled from *N* `(1`, `1)` to the individuals from the second population. We estimated a global eGRM for the long chromosome and used `GCTA` to estimate the 20 first principal components (with tag ’–pca’). We incorporated these principal components as fixed effects in `GCTA` to correct for population stratification (with tag ’–qcovar’).

### Estimating the power to find associations

#### Null simulations

In order to determine a significance cutoff for each simulation configuration, we used 300 ARG replicates from the "one population" set, and assigned each individual from each ARG a random *N* `(0`, `1)` phenotype value irrespective of genotype. We performed association tests for each association method, for each variant set / tree type and for each testing window size, i.e. for every power simulation configuration that affects the number of association tests. We set the significance cutoff such that the family-wise error rate (i.e. the fraction of replicates containing at least one significant association) was 5%.

#### Power as a function of genetic architecture

For each parameter combination of variant set / tree type, causal variant proportion, causal window size, testing window size and local heritability, we counted the number of replicates for which the *p*-value of a at least one window (for ACAT-V, local eGRM and local GRM) or variant (for GWAS) exceeded the significance threshold defined with the null simulations.

### Application to ***CREBRF***

#### Transforming the phenotypes

Phenotype data for Body Mass Index (BMI) was available for 5371 people from the Hawaiian population of the Multiethnic Cohort (MEC, Kolonel et al., 2000), along with sex and age. To ensure that the phenotype residuals would follow a standard normal distribution, we performed a transformation typical for BMI data. Namely, we stratified by sex, regressed out age and age squared, and removed individuals for which the residual was more than six standard deviations removed from the sex’s mean. Then, we inverse rank normalized the phenotypes (McCaw et al., 2020).

#### Estimating the ARG with

**`Relate`** In total, 5,384 self-identified Native Hawaiians from the Multiethnic Cohort (MEC) were genotyped on two separate GWAS arrays: Illumina MEGA and Illumina Global Diversity Array (GDA). After taking the intersection of SNPs found on both arrays, we removed variants that were genotyped in fewer than 95% of people in the sample, variants out of Hardy-Weinberg Equilibrium (*`p` <* `10`*^−^*^6^). We also applied a filter for people with more than 2% missing genotypes but removed no one with this filter.

With approximately 990,000 SNPs after quality control, we phased the genotypes with `EAGLE` (Loh et al., 2016) by using its default hg38 genetic map. We inferred ancestral alleles by using the `Relate` add-on module with ancestral genome homo_sapiens_ancestor_GRCh38_e86.tar.gz, downloaded from (ftp://ftp.ensembl.org/pub/release-86/fasta/ancestral_alleles/). We divided the genome into segments containing 10,000 SNPs, and ran `Relate` on these segments in parallel with all default parameters per the user manual.

#### Inferring the global eGRM to correct for population structure

We first inferred the segment-wise eGRMs for all chromosomes except chromosome 5, which contains the gene of interest *CREBRF*. We combined the segment-wise eGRMs into a global eGRM by taking their weighted sum, where the weights were given by the the expected number of mutations in each eGRM, which is a parameter that is provided in the output of `egrm` (Fan et al., 2022).

#### Determining the significance cutoffs

The cutoffs in Table S1 were calculated for a genomic region of length 1Mb. We compute the effective number of independent tests for each method as the number of tests for which the significance cutoff we obtain corresponds to a Bonferroni correction. For GWAS, the effective number of tests was 281.2, and for local eGRM with 5kb testing windows, 30.8. The standard genome-wide significance for GWAS is `5` *×* `10`*^−^*^8^, which corresponds to the cutoff for one million independent test with Bonferroni correction. To approximate the genome-wide cutoff value for local eGRM, we assume that the ratio of 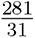 GWAS tests to local eGRM tests for a given region holds across the genome. We thus set the genome-wide local eGRM cutoff to 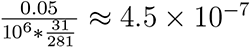).

#### Testing for QTLs

We ran our local eGRM method to test for correespondence between the transformed phenotypes and the estimated ARG around the *CREBRF* region in windows of 5kb. We corrected for population stratification by using `GCTA` to estimate the first 20 principal components of the global eGRM, leaving out chromosome 5, which we then included as fixed effects in the linear mixed model. We further used `PLINK` (Purcell et al., 2007, version 1.07) to test for association between the genotypes within the *CREBRF* region and the transformed phenotypes. To generate principal components as covariates for GWAS, we held out chromosome 5, and generated the 20 PCs using `EIGENSTRAT` (Price et al., 2006) after additionally filtering out variants with minor allele frequency < 1% and filtering for LD using commmand using –indep-pairwise 50 5 0.8 in `PLINK`.

## Results

We compare our framework based on computing the expected genetic relatedness matrix from an inferred ARG, referred to here as local eGRM, with three other association methods: GWAS, in which each variant is tested separately; local GRM, which for each testing window calculates a genetic relatedness matrix based on the typed variants within the window (see methods); and ACAT-V (Liu et al., 2019), which for each window combines the variant-level *`p`*-values from GWAS and is especially powerful when a small proportion of variants within a window are causal. Although ACAT-V is most typically applied to sequence data, our focus here is on array data, and so we apply ACAT-V to simulated array data in the main text, deferring comparisons with complete data to supplementary figures.

### The multiple testing burden is smaller for window-based association tests than for GWAS

To determine the *`p`*-value cutoffs for each method, we performed null simulations for each parameter combination of variant set or tree type and testing window size, i.e. for every simulation configuration that could lead to a different number of tests required per ARG. For each parameter combination, we simulated random phenotypes for all individuals in the sample, and we recorded the smallest *`p`*-value resulting from the tests of the simulated chromosome against the null phenotypes. We determined the significance cutoff such that the family-wise error rate was 5% in null simulations (Table S1). Fig. 2 shows the ordered *`p`*-values for one of these null simulations. It shows the general pattern that can be seen for all parameter combinations, namely that the multiple testing burden is highest for GWAS and lower for the window-based association tests ACAT-V, local GRM and local eGRM. We can compare the results in terms of the number of "effective tests" implied by the *`p`*-value cutoffs necessary to achieve a family-wise error rate (FWER) of 0.05—that is, the number of tests that would lead to the same cutoff under a Bonferroni correction. For 5kb windows and `Relate` trees, the local eGRM method applied to a 1 megabase window entails *≈* `31` effective tests. In contrast, GWAS on typed variants implies *≈* `280` effective tests, or *≈* `9` times as many as the local eGRM method. Further, the cutoffs are more stringent for GWAS when all variants are used rather than only the subset of variants selected for genotyping, whereas the difference between using only typed variants and all variants is much smaller for the window-based methods.

**Figure 2:**
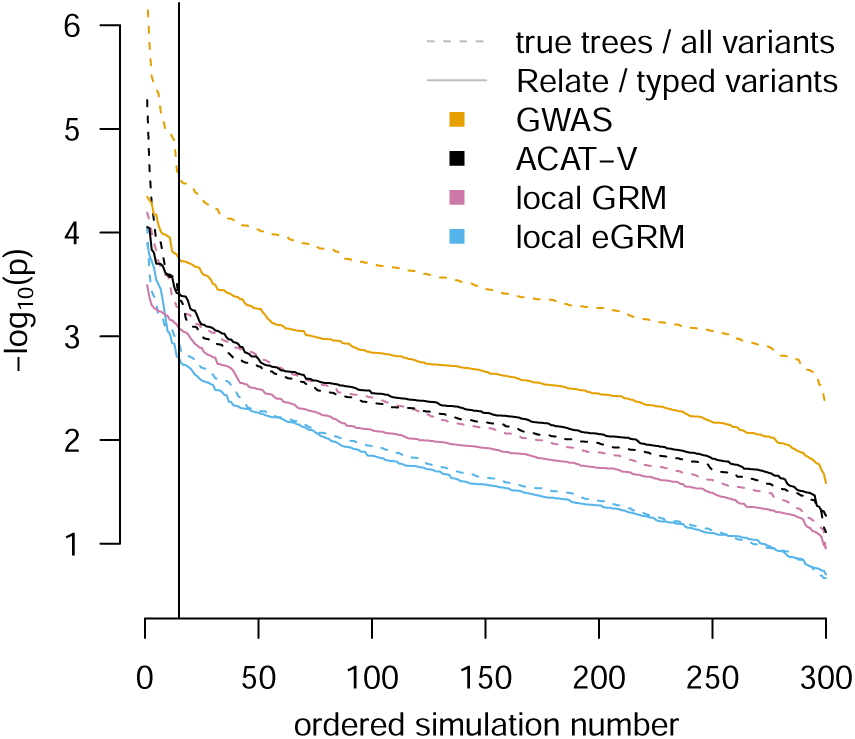
Setting *p*-value cutoffs for family-wise error rate of 5%.. The most significant *`p`*-value for any window (local GRM, local eGRM, ACAT-V) or any SNP (GWAS) in each of the 300 replicates is shown on the y-axis for each association method and ARG type / variant set, starting with the lowest minimum *`p`*-values on the left. The testing window size was 5kb. The vertical black line corresponds to 5% of the replicates.

The null simulations were also useful to determine how well the local eGRM method is calibrated with regard to the distribution of *`p`*-values under the null. The quantile-quantile plots in S1 confirm for multiple simulation configurations that both local eGRM and local GRM produce close-to-uniformly-distributed but slightly conservative *`p`*-values. Details of the *`p`*-value distribution do not influence the simulation results below, since we choose the cutoff for significance empirically based on the null simulations.

### Local eGRM exhibits power advantages in cases of allelic heterogeneity

We used simulations to understand the power of our framework to find true trait-relevant genetic regions. As with the null simulations, we simulated 200 replicates of realistic human ARGs for chromosome 1 of 1000 Africans under the out-of-Africa model using `stdpopsim` (Adrion et al., 2020). We simulated phenotypes for each individual in each ARG with a variety of architectures inside a trait-relevant genomic window by varying the number of causal variants in the window, the heritability explained by variants in the window, and the size of the testing window.

We compared the power of the following approaches: GWAS on both typed and all variants, local GRM with both typed and all variants, local eGRM with Relate trees estimated from typed variants, local eGRM with true trees, and ACAT-V with both typed and all variants.

First, we investigated power in the presence of allelic heterogeneity, i.e. multiple causal variants within close physical proximity and thus genetically linked with each other. Within a predefined causal window of the genome, each untyped variant has a given probability of being causal.

Each causal variant is given a random phenotypic effect size such that loci with lower frequency minor alleles tend to be assigned larger absolute effect sizes, as is observed in human data (see Methods for details). Fig. 3 shows power results for causal window of size 5kb, with 20% of variants causal (panels A and B, median 4 causal variants per window) or 50% of variants causal (panels C and D, median 11 causal variants per window). We also varied the local heritability, and the testing window sizes (5kb for panels A and C, 10kb for panels B and D) for the window-based tests. For the results obtained with a more extensive set of simulation parameters, including results that incorporate both typed and untyped variants, see Fig. S2.

**Figure 3:**
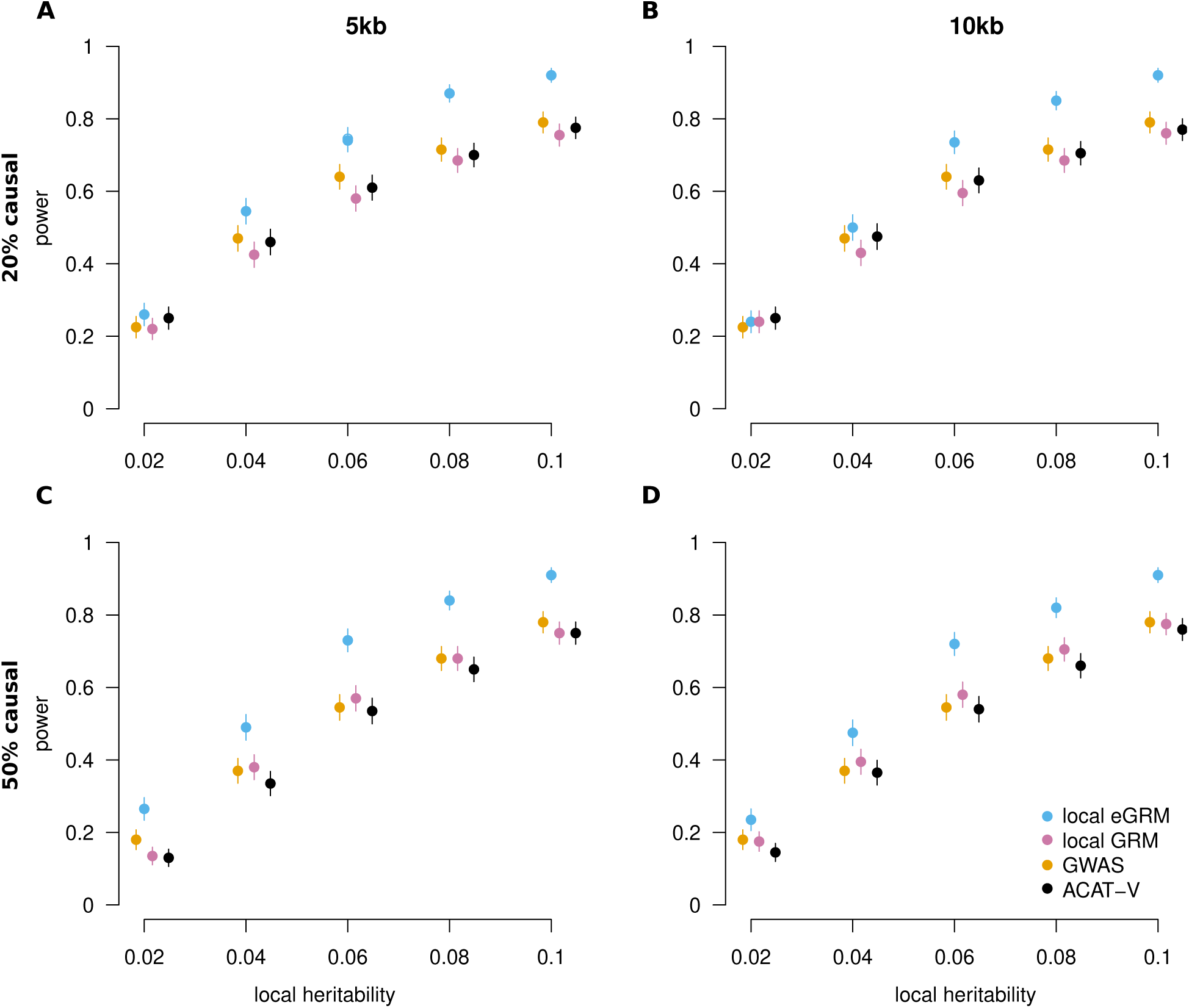
Power comparisons under allelic heterogeneity. Each panel shows the power to detect an association for 4 methods using array-like data when there is a 5kb causal window. In panels A and B, phenotypes are determined by 20% of the untyped variants in the window, while in panels C and D, phenotypes are determiend by 50% of the untyped variants in the window. In panels A and C, a 5kb test window is used (matching the simulated causal window size). In panels B and D, a 10kb test window is used. Error bars correspond to one standard error.

Across simulated genetic architectures, our local eGRM method consistently has higher power than other approaches when analyzing array data. Across the local heritability values simulated in Figure 3, the local eGRM approach with 5kb analysis windows has on average 17% higher power than GWAS with 20% of variants causal, and 31% higher power when 50% of the variants are causal. The other three methods (GWAS, local GRM, ACAT-V) performed similarly to each other. In contrast, when using the true ARG and all variant information, as would be captured by accurate sequencing data, GWAS, ACAT-V and local GRM all outperform local eGRM (Fig. S2), even when local eGRM is performed on the true trees.

Fig. 4 shows the power of each method for phenotypes that have a single untyped causal variant with allele frequency 0.02 (panel A) or 0.2 (panel B). With a single causal variant, local eGRM is roughly comparable to the other methods, operating at a slight disadvantage when the frequency of the causal variant is low (0.02), and perhaps a slight advantage when the frequency of the causal variant is higher (0.2).

**Figure 4:**
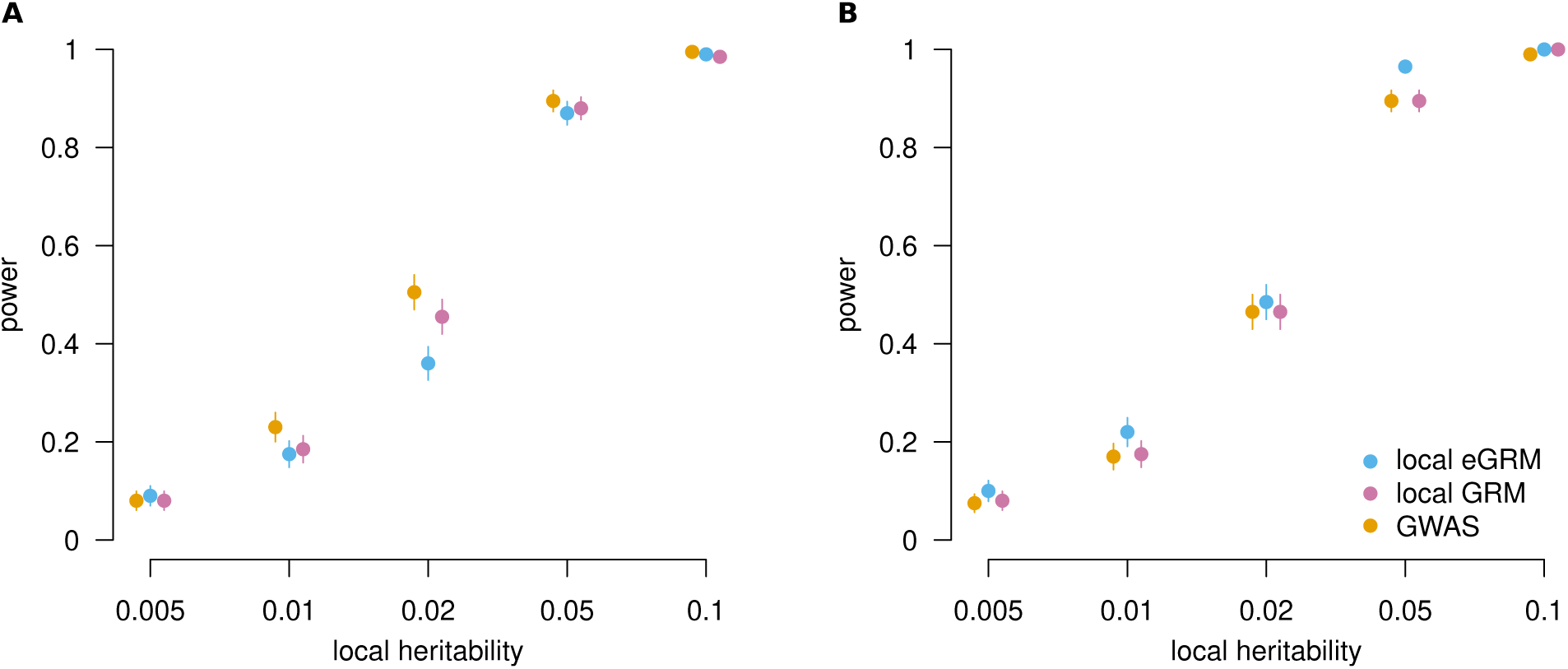
Power comparisons with one causal variant. Each panel shows the power to detect an association for 4 methods using array-like data when there is a causal variant at a frequency of either 0.02 (A) or 0.2 (B). Association tests with methods local eGRM and GRM were performed in genomic windows of 5kb. The error bars correspond to one standard error.

Tables S6 - S9 show how many replicates were found to contain a significant peak by two methods. Generally, the concordance was higher for true ARGs than for `Relate`-estimated ARGs. The highest concordance is generally between GWAS and ACAT, followed by the other pairs which have similar concordances., Tables S8 and S9 show analogous results for null simulations.

### Correcting for population stratification by including principal components inferred from the global eGRM

The simulations of the power analysis were performed on samples from a panmictic population. In real GWAS settings, however, samples are often affected by population stratification (Pritchard and Rosenberg, 1999; Rosenberg and Nordborg, 2006; Vilhjálmsson and Nordborg, 2013; Veller and Coop, 2023), in which genotypes appear correlated with phenotypes because of confounding rather than because of close linkage to causal variants. In GWAS, the most popular strategies for correcting for population stratification are inclusion of a random effect for the global GRM (Yu et al., 2006) and inclusion of fixed effects for the first several principal components of a standardized genotype matrix (Price et al., 2006), obtained by eigendecomposition of a GRM.

In order to test whether such a population stratification correction strategy works for local eGRM, we simulated a simple case of two discrete populations. In particular, we simulated a sample of 2000 diploids coming from two populations that separated 10,000 years ago. We simulated random phenotypes for all individuals, and added a fixed effect to individuals in one of the populations to simulate severe confounding due to population structure. We then computed PCs from the global eGRM derived from non-causal loci and tested a causal region for the presence of a QTL using local eGRM in windows of size 5kb, once without correcting for population stratification, and twice more correcting by including either one PC or twenty PCs as fixed effects in the model. Figure S4 shows the distribution of *`p`*-values resulting from the association tests. Panel A without the PC correction shows a very clear inflation of significant *`p`*-values—all *`p`*-values are effectively zero. Panels B and C with correction shows a *`p`*-value distribution that is very similar to the distribution of a single population sample without stratification (Fig. S1) i.e. slightly conservative but almost uniform. In this simple case, one PC should be sufficient to describe population structure (McVean, 2009), and correcting for additional PCs does not much change the distribution of *`p`*-values under the null. Thus, PC correction on a global eGRM appears able to ameliorate population stratification, at least in the simplest case. We defer a more thorough investigation of eGRM-based correction for population stratification—including investigation of more complex population structure and correction via a random-effects approach using the global eGRM—for future work.

### Analysis of a known QTL that GWAS cannot identify in the absence of a population-specific imputation panel

Lin et al. (2020) studied the limitations of imputation when there is incomplete representation of some populations in imputation reference panels. As a demonstration, they used the human locus containing gene *CREBRF*. In Pacific Islanders, there is a segregating missense mutation in *CREBRF*, rs373863828, with a large effect on adiposity. The frequency of this missense variant is as high as 26% in Samoans, but is very rare or unknown in people without recent ancestry from Polynesia. The association signal was originally detected with body mass index (BMI) in Samoans (Minster et al., 2016) at a linked tag SNP, rs12513649, which was on the Affymetrix 6.0 array. However, rs373863828 was not observed in any publicly available databases, not even those that include diverse populations (e.g. 1000 Genomes Project or Haplotype Reference Consortium), and so it was not well imputed at the time of the study. Lin et al. (2020) genotyped variant rs373863828 in self-reported Native Hawaiians who were part of the Multiethnic Cohort (MEC). When Lin et al. (2020) tested the genotyped rs373863828 directly, they found a strong association with adiposity phenotypes. However, when they used the original genotypes in the MEC, which did not contain rs373863828, they were not able to discover a significant association using 1000 Genomes Project Phase 3 and Haplotype Reference Consortium reference panels via either GWAS or admixture mapping.

Because local eGRM does not rely on an imputation panel, we explored whether local eGRM could find an association between BMI and the *CREBRF* locus in the Native Hawaiian subset of the Multiethnic Cohort (MEC, Kolonel et al., 2000) using only genotyping array data. Based on the phased genotypes, we used `Relate` to estimate the ARG for the whole genome. We then used `egrm` (Fan et al., 2022) to infer the global eGRM using the ARG of all chromosomes except chromosome 5 (the location of *CREBRF*). We then used local eGRM to test for association between the transformed phenotypes of 5371 individuals and the ARG, correcting for population stratification by including 20 PCs of the global eGRM in the linear mixed model. As can be seen in the Manhattan plot of our association results (Figure 5), we replicate Lin et al. (2020)’s inability to find a genome-wide significant association near *CREBRF* (pink shaded window) using GWAS (orange dots). However, local eGRM (blue dots) identifies signals that surpass genomewide significance in GWAS (dashed orange line) on both sides of the causal SNP rs373863828 (solid pink line). Additional windows near *CREBRF* surpass the cutoff we posit for genome-wide significance using local eGRM with 5kb testing windows based on our null simulations (blue dashed line). The distances between the observed peaks and the causal SNP rs373863828 are in line with the distances between peaks and causal variants at frequency 0.02 observed in our simulations (Table S3 and Fig. S5).

**Figure 5:**
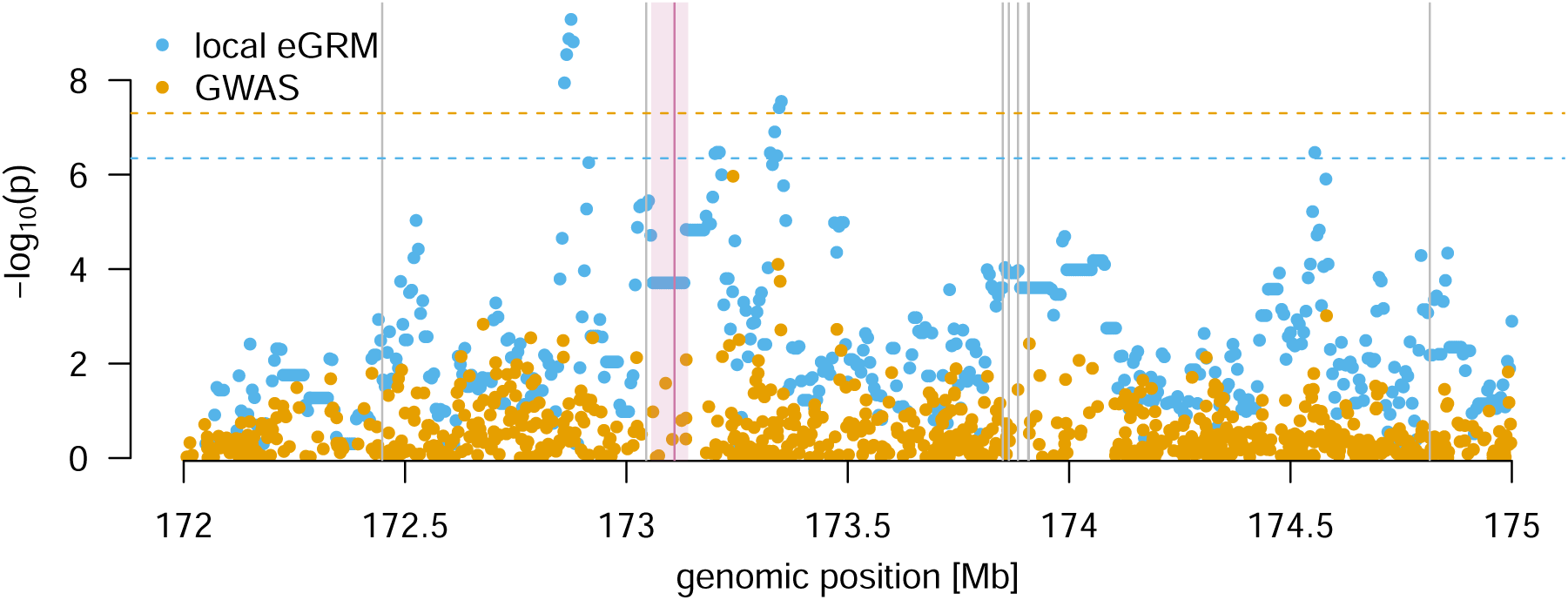
Association results for Hawaiian cohort of the MEC around *CREBRF*. Blue dots are per-window negative log_10_ *`p`*-values for the local eGRM, and orange dots are per-SNP results for GWAS. The horizontal dashed lines are the genome-wide significance cutoffs (`5` *∗* `10`*^−^*^8^ for GWAS and `4.5` *∗* `10`*^−^*^7^ for local eGRM). The vertical shaded area delimits the coordinates of *CREBRF*, and the red vertical line within it is the location of the causal SNP rs373863828. The gray vertical lines are SNPs found in the GWAS catalog that were found to be associated to traits “Body Mass Index” and “weight”. Their names, studies in which they were found, and the smallest sample size in which they were found are: rs570053489 (Tachmazidou et al., 2017, 267,616 individuals), rs12513649 (Minster et al., 2016, 3,072 individuals), rs192829047 (Zhu et al., 2020; Kichaev et al., 2019; Pulit et al., 2019, 457,822 individuals), rs3849724 (Akiyama et al., 2017; Hoffmann et al., 2018, 158,284 individuals), rs10037781 (Sakaue et al., 2021, 165,419 individuals), rs4867732 (Sakaue et al., 2021, 165,419 individuals), rs34017767 (Sakaue et al., 2021, 165,419 individuals), rs114653914 (Tachmazidou et al., 2017, 267,616 individuals). The proxy SNP rs12513649 for the *CREBRF* association is the one directly to the left of *CREBRF*.

The grey vertical lines in Figure 5 are hits for "Body Mass Index" or "weight" identified in the GWAS catalog. The one immediately to the left of *CREBRF* rs12513649, the tag SNP identified in a sample of *`∼`*3k Samoans (Minster et al., 2016). The other hits were identified in samples much larger—by factors of at least thirty—than the one we analyze here. As seen in Figure 5, local eGRM *`p`* values, though they do not usually reach genome-wide significance at this sample size (except possibly near the rightmost hit), do appear to show some degree of elevation near most of the GWAS hits.

## Discussion

We developed a new approach to QTL mapping that uses estimated ARGs to characterize local relatedness and show that it provides advantages complementary to several existing approaches to QTL mapping with SNP array data. Specifically, our approach lowers the multiple testing burden, is robust to allelic heterogeneity, and can assist in identifying QTLs even when the causal loci are not well tagged by any single array SNP and cannot be imputed because of a lack of a population-specific reference panel.

In cases of allelic heterogeneity, a marker variant can be linked with multiple causal variants that can have opposing effects, leading to their association signals interfering and causing difficulties for GWAS. The local GRM and local eGRM approaches we consider here both naturally accommodate allelic heterogeneity, because even if there are multiple causal variants in a trait-relevant region, it should still be the case that individuals who are more closely related in the region tend to be more similar on the phenotype. The local GRM and eGRM approaches differ in that the local eGRM takes into account information about local branch lengths drawn from mutations occurring in neighboring regions, since trees in neighboring regions tend to share many of the same coalescent events with the focal region. Thus, the local eGRM can capture local genetic relatedness more accurately than the local GRM, particularly for small testing windows, giving it an advantage over local GRMs formed from array data.

We identified a region surrounding the *CREBRF* gene as a QTL for BMI in a sample of Native Hawaiians where GWAS previously could not identify a genome-wide significant signal. The causal variant is not well imputed because of a lack of a population-specific imputation panel. In some sense, imputation followed by GWAS on imputed markers is conceptually roundabout: an ARG-like structure is often inferred in order to perform imputation, such as by the Li & Stephens approach (Li and Stephens, 2003), which is also the basis for recent approaches to ARG estimation (Speidel et al., 2019; Kelleher et al., 2019). In our approach, we test the structure on which imputation is performed—that is, the approximate local tree—rather than the imputed variants. Such an approach may facilitate the identification of trait-associated loci in understudied populations.

Our method adds to a long list of approaches for identifying trait-associated loci. First, and perhaps most obviously, our method is a tree-based version of methods to test local GRMs for concordance with a phenotype (Nagamine et al., 2012; Uemoto et al., 2013; Gusev et al., 2013; Wang et al., 2013; Caballero et al., 2015; Sasaki et al., 2015). As discussed above, the advantage of our approach over such methods stems from better estimates of local relatedness achieved by estimated ARGs. Further, our method can be seen as a generalization of identity-by-descent (IBD) mapping (Albrechtsen et al., 2009; Browning and Thompson, 2012; Gusev et al., 2013), where our method considers putative IBD over short regions as estimated by local trees in addition to the relatively large (multiple centiMorgan) segments that can be identified as recent IBD. IBD mapping, in turn, can be seen as a generalization of linkage mapping that uses IBD among pairs of people who are not closely related rather than only among close relatives. Our method is also closely related to haplotype mapping, and in particular approaches to haplotype mapping that estimate tree-like structures to describe relatedness among sets of haplotypes (Liu et al., 2001; Morris, 2005; Selle et al., 2021). Finally, our method adds to a tradition of methods for identifying trait-involved loci that are explicitly tree-based (Templeton et al., 1987; McPeek and Strahs, 1999; Larribe et al., 2002; Morris et al., 2002; Zöllner and Pritchard, 2005; Minichiello and Durbin, 2006; Mailund et al., 2006; Tachmazidou et al., 2007; Kimmel et al., 2008; Wu, 2008; Besenbacher et al., 2009; Zhang et al., 2012; Burkett et al., 2013; Thompson and Kubatko, 2013; Thompson et al., 2016). Whereas most previous tree-based approaches to mapping were limited to samples in the dozens because of difficulties with ARG estimation, modern ARG estimation frameworks enable a substantial gain in power using sample sizes into the thousands.

Another recent approach that used large estimated ARGs to identify trait-associated loci came from Zhang and colleagues (2021), who used a novel ARG estimation method, `ARG-Needle`, to identify trait-associated variants in a sample of over 300,000 people. Our approach is complementary to theirs. Whereas Zhang and colleagues also identify and leverage the eGRM, which they term the ARG-GRM, they use it for genome-wide tasks such as heritability estimation rather than calculating the eGRM for a local region. In their searches for trait-associated variants, they sample mutations from the ARG and test them individually, which is equivalent to testing branches or clades from the ARG. A promising future direction is to combine our approach with theirs, using our method to prioritize regions and then sampling mutations within that region in an attempt to localize the signal.

Both our results and those of Zhang and colleagues (2021) point to advantages of using estimated ARGs in situations in which genotype data are incomplete. In contrast, with complete data on underlying genetic variants, our simulations suggest that our tree-based approach is outperformed by other methods. This is sensible: in the scenarios we simulate, if all variants are known, then the tree provides no additional information. The local coalescent trees are helpful when data are incomplete because they provide a guide to the structure of unobserved mutations.^1^

Local coalescent trees could in principle outperform full sequence data in other settings as well. One such setting is in combination with a model for natural selection on trait-associated variants. Selection will distort local trees, and thus signals of selection inferred from the trees might be used to prioritize trees or clades for investigation with respect to traits that could have been under selection in the history of the sample. Another relevant setting is ascertainment, in which individuals are sampled for inclusion in the study on the basis of their trait values. Such ascertainment mimics natural selection in that it creates a sample of individuals selected on their phenotypes, and distortions in local trees under ascertainment could serve as evidence that the local region is trait-associated.

Our work here is an initial report of some advantages of a tree-based local relatedness approach to QTL mapping. The limitations of our current approach raise promising avenues for future investigation.

Here, we included all branches in the ARG within a genomic window in the eGRM, and we weighted them as a function of their branch length, span in the genome, and the number of tips descending from them. In principle, one could alter the weighting of branches, even choosing to leave some branches out, perhaps to form a time-specific eGRM (Fan et al., 2022). The absolute value of GWAS effect sizes is routinely observed to be negatively correlated with minor allele frequency, a pattern that could be explained by stabilizing selection on traits keeping large-effect variants at low frequency (Simons et al., 2018; Zeng et al., 2018; Simons et al., 2022). The "*`α`*-model" we use to simulate effect sizes is in line with the basic observation of larger effect sizes at lower-frequency variants, as is our <u>practice</u> of estimating a GRM in which variants are standardized by a factor proportional to 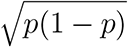, which is equivalent to assuming that the contribution to heritability of a causal variant does not depend on its frequency. However, the *`α`*-model is only a loose match to the observed distribution of effect sizes as a function of allele frequency (Simons et al., 2022; Spence et al., 2022), and using approaches to normalization or weighting of branches informed by more refined models of selection on trait-associated variation could improve performance in real data.

We did not consider errors in estimation of the ARG, instead treating marginal tree estimates from `Relate` as if they represented the true marginal trees. Figure S2 shows that using estimated trees from array data decreases power compared with using the true trees. Our main focus here is hypothesis testing, but a broader consideration of local eGRMs in attempts to estimate locally explained heritability will entail consideration of the effect of errors in ARG reconstruction on heritability estimates and their standard errors.

The variance-components model underlying our approach also assumes that in QTL windows, every branch will be associated with some normally distributed effect on the phenotype. This assumption is reasonable for QTLs with high levels of allelic heterogeneity, but it is worth exploring the application of methods that allow sparse architectures to the eGRM (Zhou et al., 2013). Further, whereas we test an additive architecture, it may be possible to modify our approach to look for QTLs that act in a dominant, recessive, or locally epistatic manner by computing modified local eGRMs (Weissbrod et al., 2016; Thompson et al., 2016; Hivert et al., 2021).

In a simple model of population structure, we showed that the false-positive rate of local eGRM QTL mapping can be controlled via including fixed effects for principal components of the global eGRM. At the same time, there are many remaining avenues to explore regarding population stratification and assortative mating, including the effect of more subtle forms of confounding on local eGRM results, performance with rare causal variants (Mathieson and McVean, 2012), the possibility of controlling for structure and relatedness via a random effect of a global eGRM, and the possibility of including PCs or random effects for modifications of the eGRM, such as time-specific eGRMs (Fan et al., 2022).

We used ARGs estimated by `Relate` (Speidel et al., 2019) for both simulated and real data. Although `tsinfer+tsdate` (Kelleher et al., 2019; Wohns et al., 2022) scales to much larger sample sizes than `Relate`, we used `Relate` because of evidence that it provides more accurate branch length estimates than `tsinfer+tsdate` (Brandt et al., 2021), which is reflected the observation of Fan and colleagues 2022 that `Relate`-based eGRMs are more accurate than those formed from `tsinfer+tsdate`. An approach to QTL mapping based on topology rather than branch length might open up application to much larger sample sizes via `tsinfer+tsdate`. `ARG-Needle` (Zhang et al., 2021), which is not yet released for general use, may also allow the procedures developed here to be used with tens or hundreds of thousands of individuals.

We tested for QTLs of size 5 kilobases or 10 kilobases. These sizes are arbitrary, but the approach of a window-based test also allows for flexibility. For example, windows could be chosen to form gene-level tests. It is likely possible to reduce the number of tests performed by adaptively choosing windows on the basis of the extent to which tree topologies change within the window. For example, in the test of *CREBRF* in Native Hawaiians, a single marginal tree spanned the entirety of the *CREBRF* gene, likely because the genotyping array included few SNPs within *CREBRF*. Testing this marginal tree only once is more sensible than testing identical windows repeatedly, as our current approach does. Building a better approach will likely require an understanding of how estimated tree topologies change as a function of sample size, population history, and the local density of typed SNPs.

Importantly, the method as currently implemented is computationally intense because of three time-consuming steps: estimating approximate ARGs with `Relate`, computing the eGRM, and fitting a linear mixed model with `GCTA`. Regarding the first step, although `Relate` is much faster than previous approaches to ARG estimation, it can still be time-consuming to run on large samples. As mentioned above, `tsinfer+tsdate` scales to larger samples than`Relate`, at the cost of less accurate branch length estimates (Brandt et al., 2021). `ARG-Needle` is reported to run on very large samples. Improvement of `tsinfer+tsdate`’s branch length estimates or release of `ARG-Needle` could allow the estimation of approximate ARGs suitable for our approach on larger samples. The second step, fitting the eGRM, is slow in very large samples because the computation entails a component for every branch on the ARG. As noted above, our approach to eGRM estimation is slower than it might be because we touch redundant branches of local trees multiple times, which can be ameliorated via a branch-based approach to computing local eGRMs (Ralph et al., 2020). Further, as noted by Zhang and colleagues 2021, it is possible to take a Monte Carlo approach to eGRM estimation, placing mutations on the ARG randomly at high rate. The GRM computed from these randomly placed mutations is an approximate eGRM that retains many of the advantages of the true eGRM. Fortunately, the third step of running the mixed model has been a major target for speedups among statistical geneticists, so we will be able to adopt existing approaches when working with larger samples (Loh et al., 2015; Runcie and Crawford, 2019).

Since before the time of Zaccheaus (Luke 19:4), people have been climbing trees to get a better view. Here, we explored a coalescent-tree-based approach to QTL mapping, showing that the expectation of the local GRM conditional on the ARG allows detection of QTLs under allelic heterogeneity or in cases in which genotype imputation is difficult. Local eGRMs are only one case of a general framework for computing ARG-based analogues of statistics typically computed on genetic variants (Ralph et al., 2020). The advantages of this general framework for a broad range of statistical- and population-genetic tasks have yet to be explored.

## Acknowledgments

We thank G. Coop, J. Felsenstein, G. Gorjanc, A. Harpak, H. Lee, M. Nordborg, P. Ralph, D. Runcie, and members of the Edge, Mooney, and Pennell labs for helpful conversations. We acknowledge support from NIH grants R35GM137758 to MDE, R01HG011646 and R35GM142783 to CWKC, R01HG012133 to NM, and F31HG012159 to BD.

## Footnotes

1 This observation is in line with a "dismal theorem" of which Joe Felsenstein has spoken publicly but not yet published. Felsenstein’s dismal theorem highlights situations in which knowledge about the evolutionary process leading to variation in trait-influencing genotypes provides no additional information about trait association if the genotypes themselves are known (Felsenstein, personal communication). It is equivalent to eq. 1 of Sen & Churchill (2001).

## Notes

### Competing Interest Statement

The authors have declared no competing interest.

