## Supplementary Tables and Figures for "Tree-based QTL mapping with expected local genetic relatedness matrices"

### Supplementary information for: Tree-based QTL mapping with expected local genetic relatedness matrices

#### Supplementary tables

**Table S1: -log10 p-value cutoffs used for power analysis**

| | data type | testing window size | method | cutoff |
|---|---|---|---|---|
| 1 | all variants | 5kb | GWAS | 4.5073 |
| 2 | all variants | 5kb | ACAT-V | 3.3467 |
| 3 | true trees | 5kb | local eGRM | 2.934 |
| 4 | all variants | 5kb | local GRM | 3.2811 |
| 5 | all variants | 10kb | GWAS | 4.5073 |
| 6 | all variants | 10kb | ACAT-V | 3.016 |
| 7 | true trees | 10kb | local eGRM | 2.7097 |
| 8 | all variants | 10kb | local GRM | 3.1375 |
| 9 | typed variants | 5kb | GWAS | 3.7475 |
| 10 | typed variants | 5kb | ACAT-V | 3.4132 |
| 11 | Relate trees | 5kb | local eGRM | 2.792 |
| 12 | typed variants | 5kb | local GRM | 3.0772 |
| 13 | typed variants | 10kb | GWAS | 3.7475 |
| 14 | typed variants | 10kb | ACAT-V | 3.1431 |
| 15 | `Relate` trees | 10kb | local eGRM | 2.7262 |
| 16 | typed variants | 10kb | local GRM | 2.8091 |

**Table S2: Number of causal variant within a 5kb causal window**

| | Min | Q25 | Median | Mean | Q75 | Max |
|---|---|---|---|---|---|---|
| proportion causal 0.1 | 1.00 | 1.00 | 2.00 | 2.25 | 3.00 | 8.00 |
| proportion causal 0.2 | 1.00 | 3.00 | 4.00 | 4.37 | 5.00 | 11.00 |
| proportion causal 0.5 | 3.00 | 8.00 | 11.00 | 10.96 | 13.00 | 22.00 |
| proportion causal 0.8 | 8.00 | 15.00 | 17.00 | 17.50 | 20.25 | 29.00 |

**Table S3: Distance between peak and causal window** The distances are calculated for power simulations between the window with the most significant p-value and the causal window for a local heritability of 0.02.

| | tree_type | causal_variants | method | mean | median | q25 | q75 |
|---|---|---|---|---|---|---|---|
| 1 | typed variants | prop causal 0.1 | GWAS | 209351 | 180594 | 59700 | 355187 |
| 2 | typed variants | prop causal 0.1 | ACAT-V | 201935 | 170000 | 35000 | 350000 |
| 3 | Relate trees | prop causal 0.1 | local eGRM | 183593 | 150000 | 40000 | 310000 |
| 4 | typed variants | prop causal 0.1 | local GRM | 204835 | 187500 | 45000 | 336250 |
| 5 | all variants | prop causal 0.1 | GWAS | 97655 | 4525 | 2592 | 158843 |
| 6 | all variants | prop causal 0.1 | ACAT-V | 121675 | 15000 | 0 | 220000 |
| 7 | true trees | prop causal 0.1 | local eGRM | 182775 | 140000 | 15000 | 335000 |
| 8 | all variants | prop causal 0.1 | local GRM | 148125 | 80000 | 0 | 295000 |
| 9 | typed variants | prop causal 0.2 | GWAS | 182621 | 147536 | 36656 | 317538 |
| 10 | typed variants | prop causal 0.2 | ACAT-V | 186250 | 152500 | 38750 | 335000 |
| 11 | Relate trees | prop causal 0.2 | local eGRM | 188050 | 155000 | 45000 | 331250 |
| 12 | typed variants | prop causal 0.2 | local GRM | 195800 | 167500 | 50000 | 340000 |
| 13 | all variants | prop causal 0.2 | GWAS | 127098 | 16242 | 2744 | 259804 |
| 14 | all variants | prop causal 0.2 | ACAT-V | 142675 | 62500 | 0 | 303750 |
| 15 | true trees | prop causal 0.2 | local eGRM | 174325 | 137500 | 15000 | 311250 |
| 16 | all variants | prop causal 0.2 | local GRM | 130725 | 67500 | 0 | 225000 |
| 17 | typed variants | causal allele frequency 0.02 | GWAS | 140091 | 92728 | 24584 | 222728 |
| 18 | typed variants | causal allele frequency 0.02 | ACAT-V | 145321 | 95782 | 28494 | 241451 |
| 19 | Relate trees | causal allele frequency 0.02 | local eGRM | 164408 | 123736 | 38857 | 254750 |
| 20 | typed variants | causal allele frequency 0.02 | local GRM | 156385 | 108843 | 32913 | 250523 |
| 21 | all variants | causal allele frequency 0.02 | GWAS | 48116 | 0 | 0 | 43896 |
| 22 | all variants | causal allele frequency 0.02 | ACAT-V | 55359 | 4694 | 2459 | 48329 |
| 23 | true trees | causal allele frequency 0.02 | local eGRM | 154987 | 102397 | 24816 | 250031 |
| 24 | all variants | causal allele frequency 0.02 | local GRM | 118146 | 50986 | 7906 | 185819 |
| 25 | typed variants | causal allele frequency 0.2 | GWAS | 104874 | 26396 | 4677 | 143892 |
| 26 | typed variants | causal allele frequency 0.2 | ACAT-V | 108341 | 27080 | 5570 | 161038 |
| 27 | Relate trees | causal allele frequency 0.2 | local eGRM | 106458 | 21426 | 6093 | 181459 |
| 28 | typed variants | causal allele frequency 0.2 | local GRM | 105460 | 16882 | 4829 | 130039 |
| 29 | all variants | causal allele frequency 0.2 | GWAS | 51896 | 0 | 0 | 10537 |
| 30 | all variants | causal allele frequency 0.2 | ACAT-V | 45782 | 2841 | 1382 | 6944 |
| 31 | true trees | causal allele frequency 0.2 | local eGRM | 59582 | 5237 | 2174 | 23863 |
| 32 | all variants | causal allele frequency 0.2 | local GRM | 82919 | 14220 | 2272 | 90098 |

**Table S4: Proportion of replicates with shared significant hits - one causal variant** Results are for `Relate` trees / typed variants, causal variant frequency 0.2, local heritability 0.02 and testing window size 5kb

| | GWAS | ACAT | local eGRM | local GRM |
|---|---|---|---|---|
| GWAS | 0.47 | 0.45 | 0.35 | 0.41 |
| ACAT | | 0.47 | 0.36 | 0.41 |
| local eGRM | | | 0.48 | 0.36 |
| local GRM | | | | 0.47 |

**Table S5: Proportion of replicates with shared significant hits - one causal variant** Results are for true trees / all variants, causal variant frequency 0.2, local heritability 0.02 and testing window size 5kb

| | GWAS | ACAT | local eGRM | local GRM |
|---|---|---|---|---|
| GWAS | 0.73 | 0.68 | 0.56 | 0.50 |
| ACAT | | 0.69 | 0.55 | 0.49 |
| local eGRM | | | 0.59 | 0.47 |
| local GRM | | | | 0.52 |

**Table S6: Proportion of replicates with shared significant hits - one causal window** Results are for `Relate` trees / typed variants, causal window size 5kb, proportion of causal variants 0.1, local heritability 0.02 and testing window size 5kb

| | GWAS | ACAT | local eGRM | local GRM |
|---|---|---|---|---|
| GWAS | 0.24 | 0.20 | 0.18 | 0.17 |
| ACAT | | 0.23 | 0.17 | 0.17 |
| local eGRM | | | 0.27 | 0.15 |
| local GRM | | | | 0.21 |

**Table S7: Proportion of replicates with shared significant hits - one causal window** Results are for true trees / all variants, causal window size 5kb, proportion of causal variants 0.1, local heritability 0.02 and testing window size 5kb

| | GWAS | ACAT | local eGRM | local GRM |
|---|---|---|---|---|
| GWAS | 0.50 | 0.45 | 0.20 | 0.29 |
| ACAT | | 0.45 | 0.18 | 0.27 |
| local eGRM | | | 0.27 | 0.20 |
| local GRM | | | | 0.36 |

**Table S8: Proportion of replicates with shared significant hits - null simulations** Results are for `Relate` trees / typed variants, random phenotypes with no causal variants and testing window size 5kb

| | GWAS | ACAT | local eGRM | local GRM |
|---|---|---|---|---|
| GWAS | 0.05 | 0.03 | 0.02 | 0.02 |
| ACAT | | 0.05 | 0.01 | 0.02 |
| local eGRM | | | 0.05 | 0.01 |
| local GRM | | | | 0.05 |

**Table S9: Proportion of replicates with shared significant hits - null simulations** Results are for true trees / all variants, random phenotypes with no causal variants and testing window size 5kb

| | GWAS | ACAT | local eGRM | local GRM |
|---|---|---|---|---|
| GWAS | 0.05 | 0.04 | 0.02 | 0.02 |
| ACAT | | 0.05 | 0.02 | 0.02 |
| local eGRM | | | 0.05 | 0.02 |
| local GRM | | | | 0.05 |

**Table S10: Association of PCA loadings and Body Mass Index** Linear regression results for the transformed BMI and the first 50 eigenvectors of the genome-wide eGRM (except chromosome 5).

| | log10(p) | R-squared | slope |
|---|---|---|---|
| PC1 | -0.9068 | 0.0004 | -1.5323 |
| PC2 | -2.1875 | 0.0014 | 2.7139 |
| PC3 | -0.2788 | 0.0001 | -0.6310 |
| PC4 | -5.5871 | 0.0041 | -4.6902 |
| PC5 | -0.0171 | 0.0000 | -0.0482 |
| PC6 | -0.4491 | 0.0002 | 0.9220 |
| PC7 | -1.6967 | 0.0010 | -2.3141 |
| PC8 | -4.4827 | 0.0032 | -4.1321 |
| PC9 | -2.4156 | 0.0016 | 2.8789 |
| PC10 | -0.3385 | 0.0001 | 0.7379 |
| PC11 | -5.8140 | 0.0043 | -4.7859 |
| PC12 | -25.8888 | 0.0211 | -10.5891 |
| PC13 | -0.0028 | 0.0000 | 0.0079 |
| PC14 | -1.1218 | 0.0006 | 1.7704 |
| PC15 | -1.1247 | 0.0006 | 1.7730 |
| PC16 | -0.2384 | 0.0001 | -0.5546 |
| PC17 | -3.4650 | 0.0024 | 3.5674 |
| PC18 | -0.0029 | 0.0000 | 0.0084 |
| PC19 | -1.0501 | 0.0005 | -1.6945 |
| PC20 | -0.2870 | 0.0001 | 0.6467 |
| PC21 | -0.1281 | 0.0000 | -0.3246 |
| PC22 | -1.6011 | 0.0009 | -2.2320 |
| PC23 | -0.2698 | 0.0001 | -0.6146 |
| PC24 | -0.4274 | 0.0001 | -0.8862 |
| PC25 | -1.0185 | 0.0005 | 1.6607 |
| PC26 | -0.6581 | 0.0003 | -1.2227 |
| PC27 | -0.2063 | 0.0000 | -0.4914 |
| PC28 | -0.6813 | 0.0003 | -1.2540 |
| PC29 | -0.5015 | 0.0002 | -1.0005 |
| PC30 | -0.0741 | 0.0000 | 0.1974 |
| PC31 | -1.6213 | 0.0010 | -2.2501 |
| PC32 | -1.7001 | 0.0010 | -2.3230 |
| PC33 | -0.1268 | 0.0000 | -0.3218 |
| PC34 | -0.0957 | 0.0000 | -0.2496 |
| PC35 | -2.6210 | 0.0017 | -3.0315 |
| PC36 | -1.3555 | 0.0008 | -2.0058 |
| PC37 | -1.1639 | 0.0006 | 1.8146 |
| PC38 | -0.0204 | 0.0000 | -0.0574 |
| PC39 | -0.2452 | 0.0001 | 0.5681 |
| PC40 | -0.0376 | 0.0000 | 0.1037 |
| PC41 | -1.1126 | 0.0006 | -1.7604 |
| PC42 | -0.1046 | 0.0000 | 0.2706 |
| PC43 | -0.2411 | 0.0001 | 0.5601 |
| PC44 | -2.8898 | 0.0019 | -3.2065 |
| PC45 | -0.5372 | 0.0002 | 1.0538 |
| PC46 | -0.9868 | 0.0005 | 1.6270 |
| PC47 | -0.7248 | 0.0003 | 1.3099 |
| PC48 | -0.9924 | 0.0005 | -1.6309 |
| PC49 | -0.2039 | 0.0000 | -0.4881 |
| PC50 | -1.7766 | 0.0011 | -2.4042 |

#### Supplementary figures

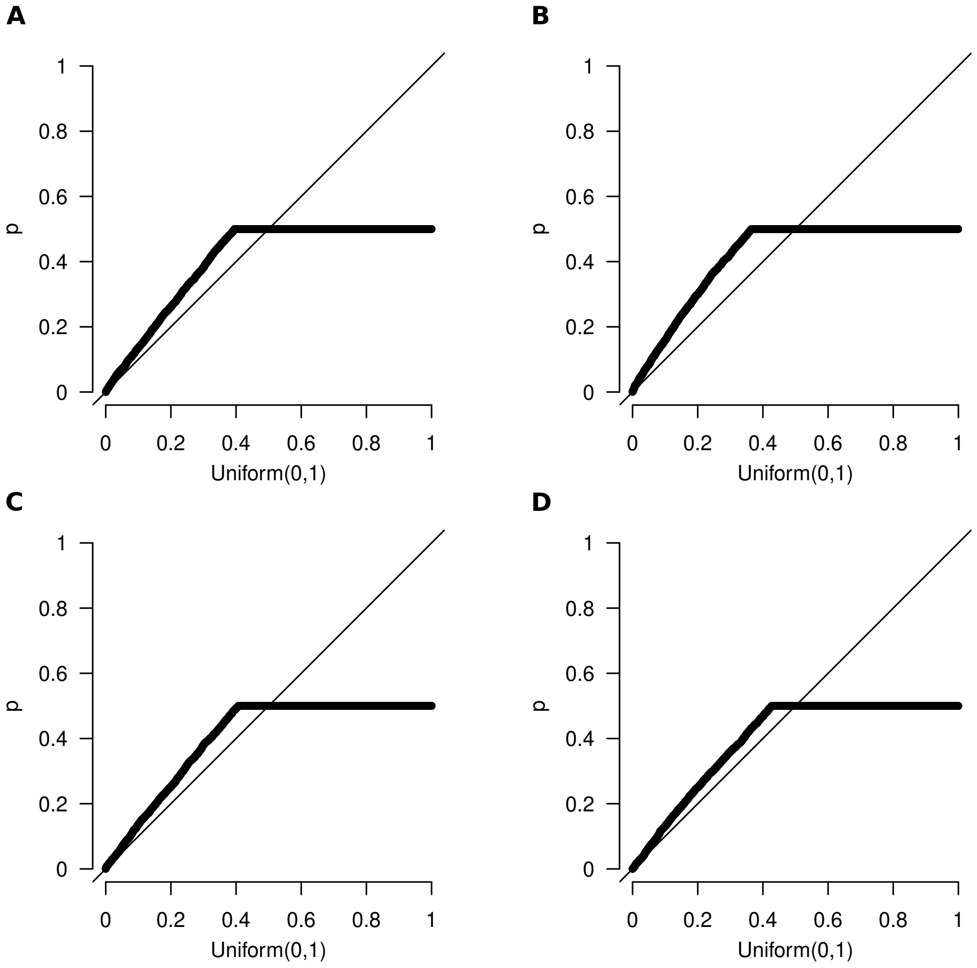


**Figure S1: Null simulations** Association results between phenotypes chosen randomly from $N(0,1)$ and an ARG of 1000 humans (extract of chromosome 1) simulated with `stdpopsim`. `Relate` trees were inferred from the genotypes of 20% of the common variants (frequency $\geq 0.01$). Association tests were performed for the eGRMs and GRMs calculated in genomic windows of 5kb. Panels A and B are results for `Relate` trees / typed variants and panel C and D for the true trees / all variants. Panels A and C are results for local eGRM and panels B and D for local GRM.

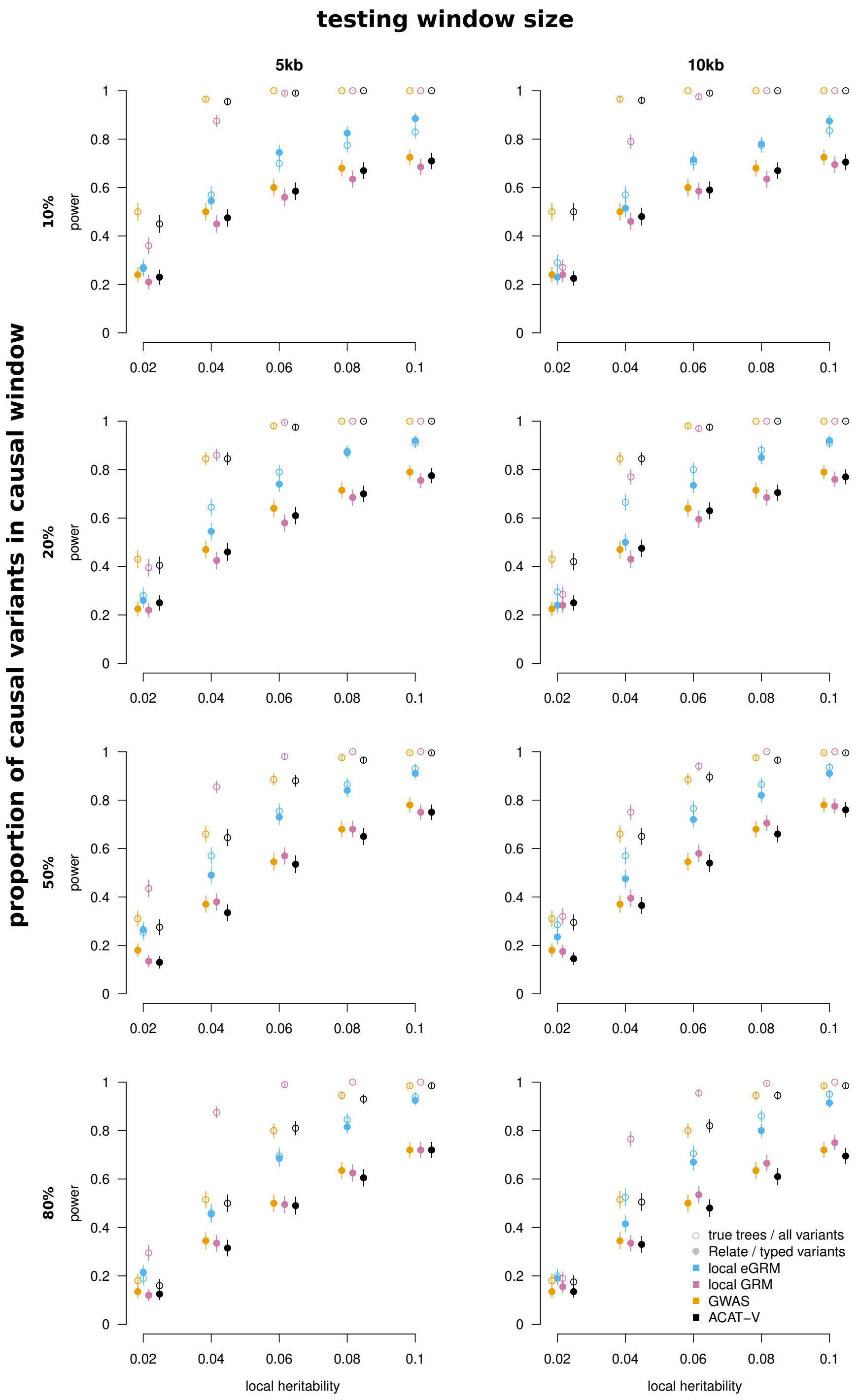


**Figure S2: Power allelic heterogeneity** Power to find a significant hit for simulated ARGs and phenotypes with one causal window of 5kb containing a certain proportion of causal variants in 200 replicates. The causal variants make up 10-80% of the untyped variants in the causal window. Association tests were performed for genomic windows of 5 and 10kb, as indicated by vertical columns. The error bars correspond to one standard error.

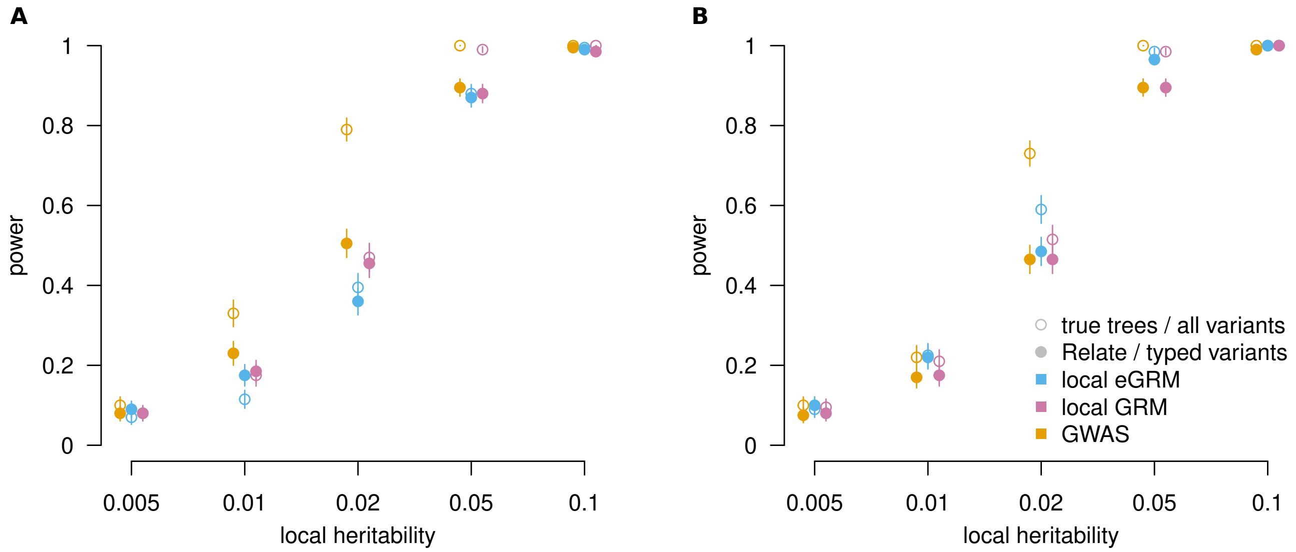


**Figure S3: Power with one causal variant** Power to find a significant hit for simulated ARG and phenotypes with one causal variant with allele frequency 0.02 (A) or 0.2 (B). Association tests with methods local eGRM and GRM were performed in genomic windows of 5kb. The error bars correspond to one standard error.

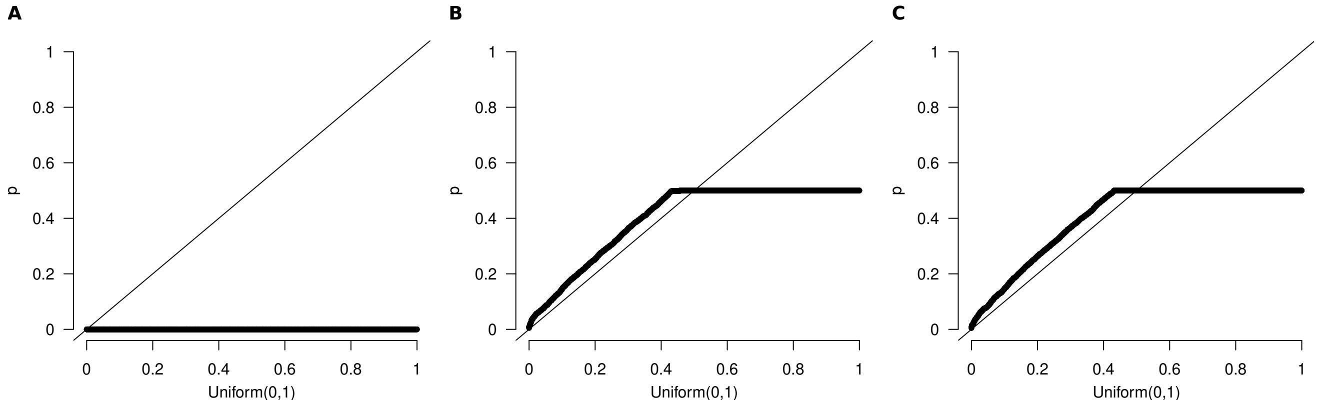


**Figure S4: Population structure** Distribution of p-values resulting from local eGRM association tests of random phenotypes with simulated ARGs consisting of 1000 diploid samples from each of two populations that split 10,000 years ago. Population stratification was achieved by adding 1 to the phenotypes of the individuals of one of the populations. (A) $p$-values of local eGRM association tests that did not correct for stratification. (B) $p$-values of local eGRM association tests that used one principal component of a global eGRM to correct for population stratification. (C) $p$-values of local eGRM association tests that used 20 principal components of a global eGRM to correct for population stratification.

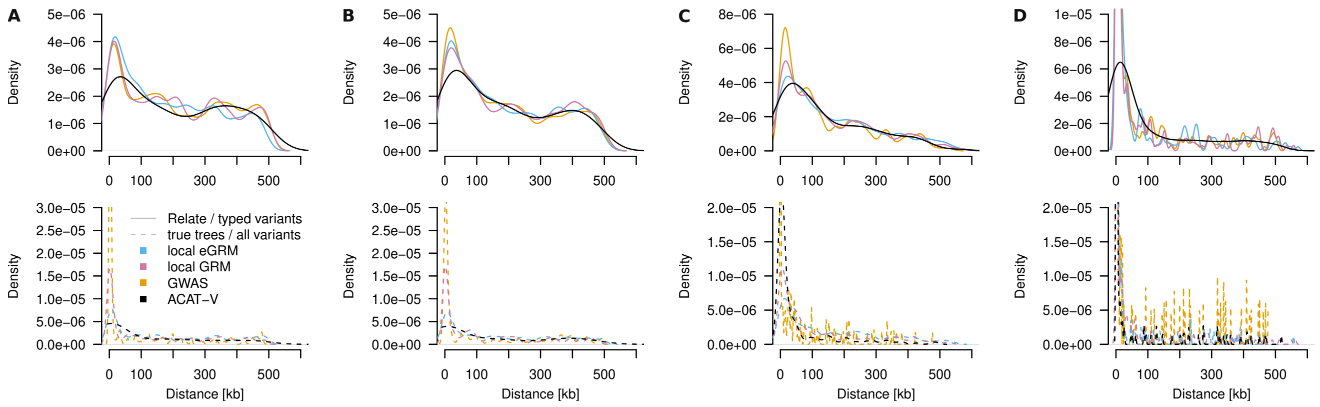


**Figure S5: Distance between peak and causal window** The distances are calculated for power simulations between the window with the most significant *p*-value and the causal window for a local heritability of 0.02 and testing window size 5kb. (A) Allelic heterogeneity simulations with causal window size 5k and proportion of causal variants 0.1. (B) Allelic heterogeneity simulations with causal window size 5k and proportion of causal variants 0.2. (C) One causal variant simulations with causal allele frequency 0.02. (D) One causal variant simulations with causal allele frequency 0.2. The density bandwidth was chosen with method "SJ" (Sheather and Jones, 1991).

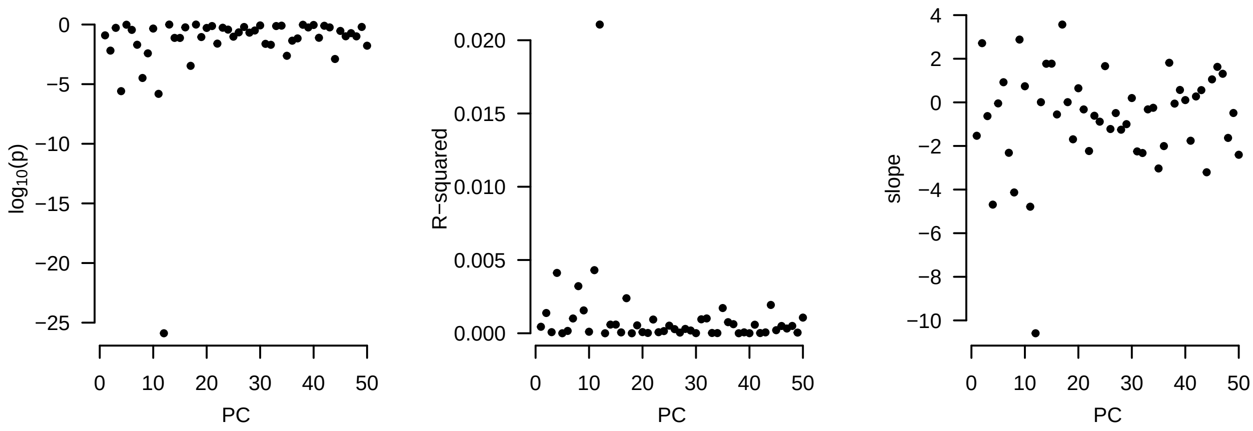


**Figure S6: Association of PCA loadings and Body Mass Index** Linear regression results for the transformed BMI and the first 50 eigenvectors of the genome-wide eGRM (except chromosome 5).
